# Converting Microwave Ovens into Plasma-Generating Decontamination Units for N-95 Respirators

**DOI:** 10.1101/2020.09.15.297630

**Authors:** David N. Ruzic, Chamteut Oh, Joseph V. Puthussery, Dhruval Patel, Zach Jeckell, Vishal Verma, Thanh H. Nguyen

**Affiliations:** University of Illinois at Urbana-Champaign, Urbana, IL, USA

**Keywords:** COVID-19, N-95 Respirator, Decontamination, Plasma, Microwaves

## Abstract

We show how a common microwave oven, a coat-hanger and a coffee cup can be used to decontaminate N-95 respirators in 30 seconds. Tulane virus in the artificial saliva was reduced by >3 log and *Geobacillus stearothermophilus* spores were reduced by >6 log. Respirators maintained filtration and fit even after 10 cycles. Filtration and fit tests performed by the CDC confirmed there was no damage to the respirators. Spectroscopy of the plasma reveals that OH and C-containing radicals as the most prevalent active species.

## I. Introduction

The ongoing pandemic of SARS-CoV-2 (i.e. COVID-19) has severely stressed the worldwide healthcare system and has created unprecedented shortages of personal protective equipment (PPE) including N-95 filtering respirators (N-95s). Plasma sterilization is a known technique and works on viruses [1]. A factor of 30 reduction of human norovirus was obtained after a 5-minute treatment from a cold atmospheric-pressure air plasma by a FlatPlasTer 2.0™ device. The inactivation of the virus particles’ functions occur through synergistic effects of the cold plasma-initiated air chemistry, which consists of nitric oxide (NO) (including its intermediates, NO radicals, NO^−^, and NO^+^, and adducts, NO_2_, NO_2_^−^, NO_3_, N_2_O_3_, N_2_O_4_, and ONOO^−^) and reactive oxygen species (including ozone, atomic oxygen, singlet oxygen, and oxygen ions).[1] In addition UV light was generated by this device. The main UV components of the emission spectrum (measured with an Avantes spectrometer, AvaSpec-2048) were in the wavelength range between 280 and 400 nm, which can be mainly attributed to the excitation of molecular nitrogen in air. That means, the UVC light was almost absent, whereas UVA and UVB were active plasma components.[2]

Commercial plasma sterilization devices, such as the FlatPlasTer 2.0™, are not widely available and are costly. Few if any hospitals, nursing homes, or medical facilities have such a device. However, everyone has a microwave oven – often on every floor or even every single room. Microwave ovens have been used to make plasmas. There are numerous how-to videos using grapes on the internet [3] and scholarly articles explaining how they work [4]. Our lab specializes in using microwave sources to make atmospheric-pressure plasmas too for a variety of uses [5]. The new idea described in this work is to create a plasma by making an antenna that has a length equal to some multiples of the wavelength of the microwaves in a microwave oven (some multiple of 12.2 cm) and bending it into a split ring with a small (2mm) gap, an intense electric field is generated in the gap. That electric field can ignite a plasma under the right conditions. Those conditions are created by adding a conducting liquid, such as saline solution, into the gap, and placing the antenna cap on a piece of ceramic. The conducting fluid electrically shorts the gap causing a large current to flow and heats the fluid quickly. The water is vaporized in a few seconds and a spark is generated in the gap. These first electrons then allow a plasma to be generated in the gap out of the air and other gasses and materials present. This plasma is sustained as long as the microwave power is present. Plasmas generally generate UV light, but UV light alone is insufficient. Plasmas also produce radicals, particularly O radicals from the air and the water, and carbon-containing radicals from the ceramic surface. Hydrogen peroxide (30%) is also added to the gap with the saline. This produces hydrogen peroxide vapor and OH radicals. Positive and negative ions and electrons are generated as well, which likely leads to ozone and NO compounds. All of these species are biologically active and will react away any mucus/saliva barriers and then destroy the cell walls of the biological contaminant.

The ability to turn a microwave oven into a decontamination device with common everyday materials such as a coat-hanger, coffee cup and some fluids means instant world-wide accessibility. Hospitals, nursing homes, meat-packing plants, etc., could practice this technique with materials at hand and perform decontaminations in as little as 30 seconds. The procedure to do this process is as follows: 1) Obtain a piece of heavy gauge wire (8 to 12 gage) which is stripped of insulation, such as a metal coat hanger. Cut it to 24.4 cm. Bend it into a circle leaving a gap of around 2mm. You have made a split-ring antenna with a length of 2 times the wavelength of the 2.45 GHz microwaves which power microwave ovens. Through experimentation the length must be within 5% ie (24 to 25 cm). The gap must be between 1 and 2.5 mm. 2) Disable the rotation in the microwave oven. This can be simply done by turning the turn-table dish upside down. 3) Place the antenna with the gap down into a glass or ceramic shallow dish. An inverted coffee cup works well. Both ends of the antenna forming the gap should be touching the ceramic/glass surface. It may need to be tilted up and supported to do that. See Figure 1. 4) Add some saline solution (around 1 ml, the exact quantity not critical) and some 30% hydrogen peroxide (at least 1ml or 1 tsp) in the antenna gap. 5) Turn the microwave on its highest power (normal operation) for 1 minute maximum. At first the liquid will start boiling. In about 15 seconds a popping sound and bright sparks will emanate from the gap. (Figure 1, lower inset) Soon, a continuous plasma will form, and a loud buzzing sound will be heard. The continuous plasma should run for at least 30 seconds. Do not leave the mask in for more than 1 minute. If the mask is left in too long (about 2 minutes) the mask material around the metal nose piece may start to melt. A how-do video [6] is available which shows the complete process and includes additional safety tips.

**Figure 1:**
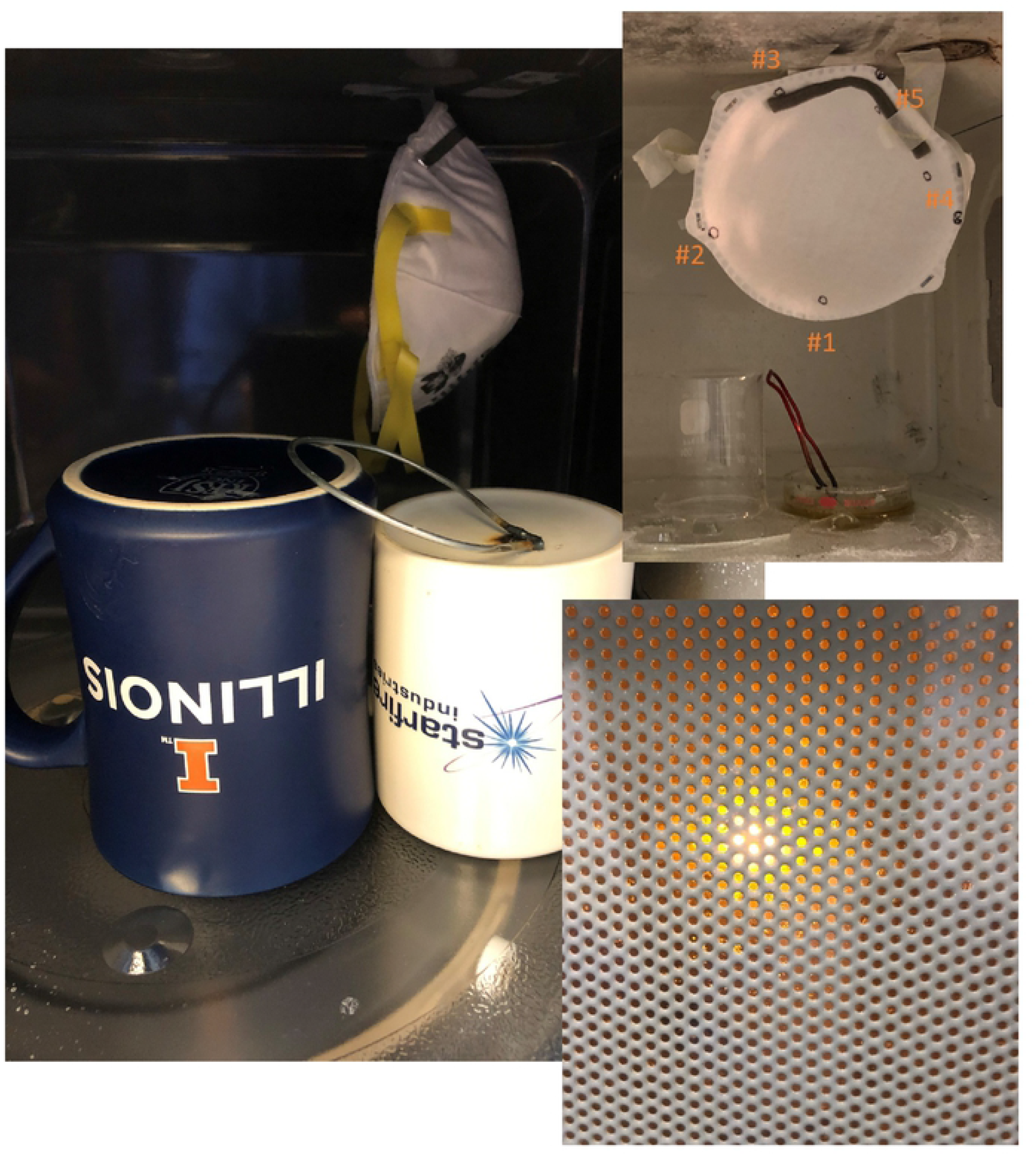
Antenna on inverted coffee cup in microwave oven with N 95 respirator suspended behind it. Upper inset shows the locations of the viruses on the mask on one side. Lowerinset is a picture of the intense plasma created at the antenna gap on the cup surface.

## II. Methods

### A. Spore inactivation experiments

*Geobacillus stearothermophilus* spore kits (EZS/5, Mesalabs) were used to determine the sporicidal efficacy of microwave plasma. The kit contained the bacterial spores (2×10^6^ CFU) on a piece of paper with culture medium in a separate glass ampule. The filter-paper was taken out of the kit and hung from the top of the microwave with three different distances (10, 20, and 30 cm) from the source of plasma (i.e., the antenna). After the decontamination treatment, the filter-paper was incubated at 60°C for 24-hour with the culture medium. The sporicidal efficacy of >6log_10_ reduction of the experimental conditions was confirmed by the violet color being unchanged. Once the violet color of the culture medium changed to yellow, this meant the >6 log_10_ reduction of spores was not achieved.

### B. Virus inactivation experiment

We used Tulane virus, which is in the calicivirus family and a surrogate for human norovirus, for testing. This virus is a mammalian virus within the same Baltimore classification as SARS-CoV-2 (i.e., (+)ssRNA). Tulane virus was received from Cincinnati Children’s Hospital Medical Center. The MA104 cells were used as the host cells in virus propagation and plaque assay as described previously [7, 8]. The testing solution was prepared by mixing 15 uL of tulane virus and 15 uL of the artificial saliva (ASTM E2720-16 with a slight modification) for two types of experiment. First, the testing solution was inoculated to respirator pieces (10 mm diameter) to determine the inactivation efficacy of the microwave oven and hydrogen peroxide vapor without the plasma generation. The respirator pieces were put in the microwave oven and exposed to 1) the microwave for 1 min without hydrogen peroxide and the antenna, 2) the microwave for 1 min with hydrogen peroxide, and 3) the microwave for 1 min with hydrogen peroxide and the antenna. Second, the testing solution was seeded to the five places inside and five places outside of the N95 respirator (Fig. 1 upper inset) to check the impact of respirator shape to the inactivation efficacy. The contaminated respirator was placed above the antenna inside the microwave oven as shown in Figure 1. The microwave oven was then turned on for 30 s with hydrogen peroxide and the antenna. After the decontamination process, each inoculation site of the respirator was cut into 10 mm diameter pieces which were submerged each in 1 mL of culture medium without FBS. The viruses were then detached from the respirator pieces by vortexing them for 3 min and shaking them for 30 min at 450 rpm. The same procedure was carried out with the negative controls which were left in the biosafety cabinet throughout the experiments (about an hour).

### C. Illinois filtration and fit testing

Circular sections of the test respirator (2 circular sections of 47 mm diameter were cut out from each respirator) were loaded on to a filter holder (URG, Carrboro, NC, USA) and exposed to dried and charge-neutralized polydisperse 2% NaCl aerosols generated using a constant output atomizer (TSI Model 3076, MN, USA). The face velocity on the filter was maintained at 9.4 cm/s, which is equivalent to an effective flow rate of 85 lpm (NIOSH recommended most severe breathing flow rate) through the entire respirator area [surface area of the respirator was measured manually (~ 150 sq cm)]. NaCl particle number concentration was measured before and after connecting the test filter using a condensation particle counter (CPC, TSI Model 3022A; flow rate = 1.5 lpm). The particle filtration efficiency of the tested filter material was calculated using the equation below:

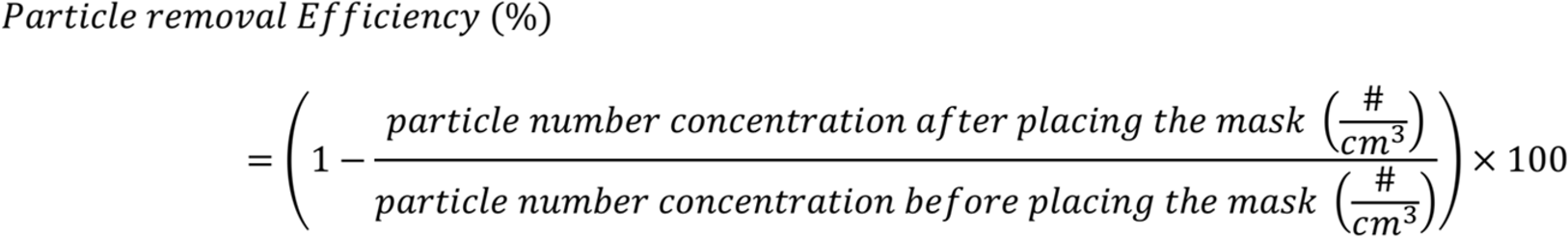

The filtration efficiency tests were performed twice on each of the respirator after 0,1, 3, 5 and 10 cycles of plasma exposure. Respirators exposed to microwave only, boiling 30% hydrogen peroxide and saline only were used as the controls.

The quantitative fit testing was performed at the University of Illinois following the modified ambient aerosol condensation nuclei counter quantitative fit testing protocol (1910.134 App A, OSHA). The N95 respirators which underwent 1, 3, 5, and 10 cycles of decontamination treatment were prepared.

### D. CDC filtration and fit testing

A set of masks were also sent to the National Personal Protective Technology Laboratory run by the CDC for evaluation. Eleven respirators (3M model 1860) that were unworn and not subjected to any pathogenic microorganisms were submitted for evaluation. This included 7 respirators that were subjected to 3 cycles of the microwave plasma decontamination process and an additional 4 respirators that served as controls. The samples were tested using a modified version of the NIOSH Standard Test Procedure (STP) TEB-APR-STP-0059 to determine particulate filtration efficiency. The TSI, Inc. model 8130 using sodium chloride aerosol was used for the filtration evaluation. For the laboratory fit evaluation, a static manikin headform was used to quantify changes in manikin fit factor. The TSI, Inc. PortaCount® PRO+ 8038 in “N95 Enabled” mode was used for this evaluation. Additionally, tensile strength testing of the straps was performed to determine changes in strap integrity. The Instron® 5943 Tensile Tester was used for this evaluation.

### E. Plasma Spectroscopy

Spectra were obtained from the plasma in the microwave oven by drilling a hole through the back of the oven and inserting an optical fiber. The fiber was connected to an Oceanoptics HR 2000+ ER Spectrometer and data between 200-1100 nm was taken at various intervals during the process. All decontamination experiments were conducted with a steel antenna made from a coat hanger, but to understand the origin of the spectral lies, additional spectra were taken when using an antenna made from Cu and Al.

## III. Results

The plasma generated by the microwave oven was effective to decontaminate both the spores and Tulane virus. As shown in Table 1, the spore tests indicated that exposure to the plasma for 30 s inactivated the spores by >6-log_10_ even in the far corner of the microwave oven (i.e., 30 cm away from the antenna). In the cases where the spore paper was hung in the same place, but a plasma was not generated (ie microwave’s only), the 6-log_10_ reduction was not achieved.

**Table 1.** Summary of the microwave oven experiments with different experiment conditions.

| Conditions | Spore test | Virus log<sub>10</sub> reduction |  | Filtration efficiency (%) | Fit score |
| --- | --- | --- | --- | --- | --- |
|  |  | Inside respirator | Outside respirator |  |  |
| Control | NA<sup>a</sup> | 0 | 0 | 99.4 | NA |
| Microwave (60 s) | Fail | 0.29±0.12<sup>b</sup> | -0.08±0.17<sup>b</sup> | 99.6 | NA |
| Microwave with H<sub>2</sub>O<sub>2</sub> and saline (60 s) | Fail | 0.9<sup>c</sup> | 4.08<sup>c,e</sup> | 99.5 | NA |
| Plasma (30 s) | Pass | 4.22±0.40<sup>d,e</sup> | 4.22±0.79<sup>d,e</sup> | 99.5 | 115±23 |

a) Not applicable.
b) Microwave alone was applied to 5 mm diameter coupons of respirator (N=3).
c) Microwave with H<sub>2</sub>O<sub>2</sub> and saline experiment were done with the 5 mm diameter coupons of respirator (N=1).
d) Plasma treatment was performed with the whole N95 respirator with virus inoculation on different locations of the respirator (N=5).
e) The actual inactivation efficacy could be higher than the presented value because of the detection limit. The detection limit of inactivation efficacy was determined by the ratio of initial viral infectivity to the viral infectivity after treatment. Initial virus concentrations were little different and thus the detection limit was also different.

Tulane viruses inoculated both sides of the respirator were reduced by a factor of greater than 3-log_10_ regardless of the inoculation sites. Neither microwave alone nor the microwave with hydrogen peroxide were able to inactivate greater than 3-log_10_ reduction, which means the plasma generation was essential for the decontamination. Although the microwave and hydrogen peroxide are also known to inactivate the viruses, the treatment time seemed too short to attain the goal of 3-log_10_ reduction.

The particle filtration efficiency of the respirator after multiple plasma exposure cycles are shown in Figure 2. The initial particle filtration efficiency of the respirators before and after plasma exposure were very similar (i.e. ≥ 99 %). Furthermore, all the tested respirators passed the fit test after 1, 3, 5 and 10 plasma exposure cycles. Collectively these results suggest that the decontamination of the N95 respirator by plasma exposure does not compromise the filtration efficiency or the fit of the N95 respirator. The filtration and fit tests done by the CDC’s NPPTP lab confirmed these results. Tables 2, 3 and 4, show that there is no statistical difference between the masks which underwent 3 cycles of decontamination compared to the control samples.

**Figure 2.**
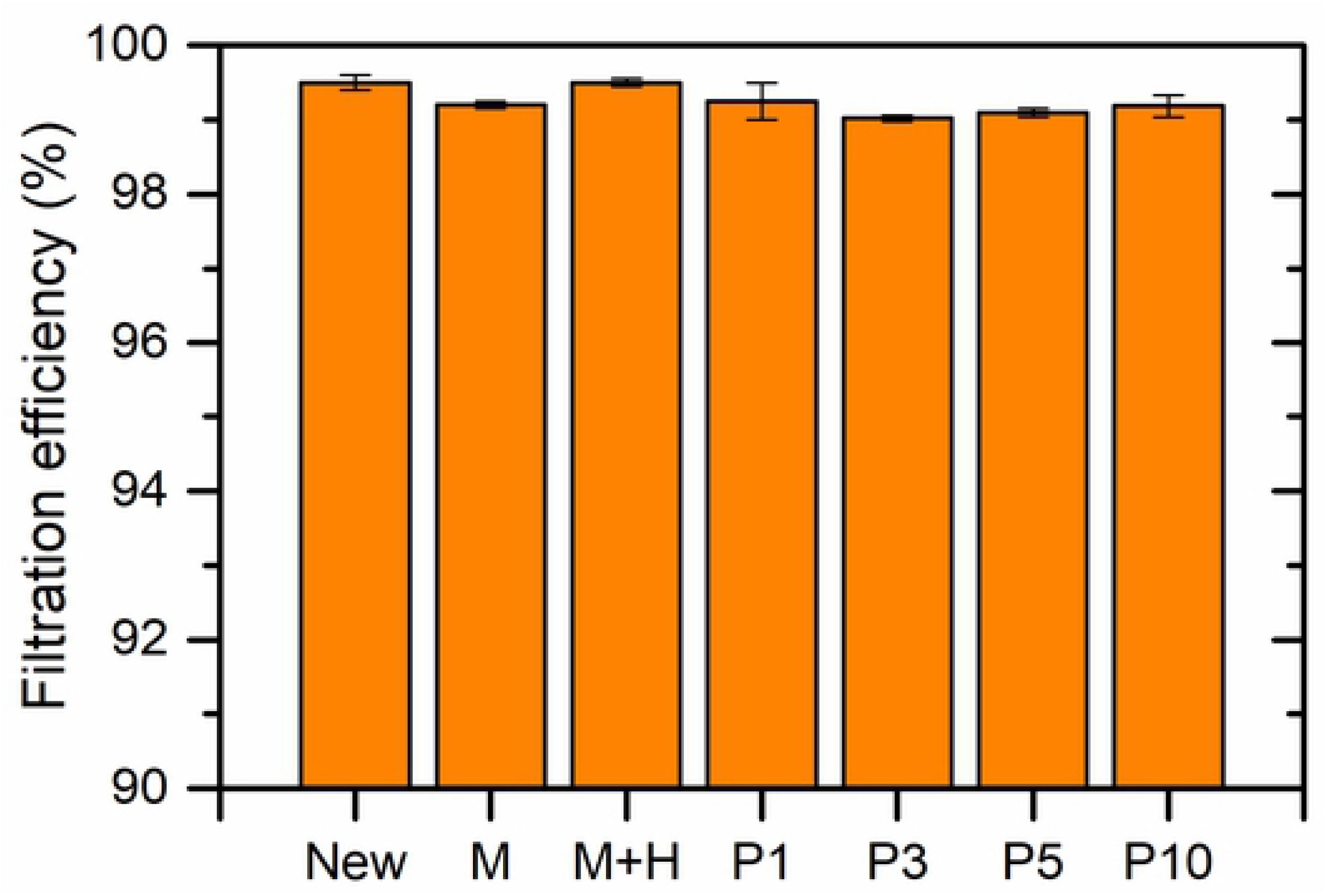
Filtration test results showing no statistical difference in filtration ability from the microwave, plasma or vapor exposures. ‘New’ stands for clean respirator before decontamination. M and H are the assumed controls after 2 minutes microwave exposure alone, and 2 minutes microwave in the presence of hydrogen peroxide and saline, respectively. Pl, P3, PS and PIO denotes the respirator after I, 3, 5 and 10 plasma decontamination cycles. The error bars denote the standard deviation (Iσ) of the duplicate measurements.

**Table 2.** Filter Efficiency Evaluationfrom NPPTP.

| Respirator Model,<br>Decon Method, #<br>of cycles | Treated<br>Sample # | Flow Rate<br>(Lpm) | Initial Filter<br>Resistance<br>(mmH<sub>2</sub>O) | Initial Percent<br>Leakage (%) | Maximum<br>Percent<br>Leakage (%) | Filter<br>Efficiency<br>(%) |
| --- | --- | --- | --- | --- | --- | --- |
| **3M 1860,**<br>**microwave**<br>**plasma, 3 cycles**<br>Min Fil Eff: 98.81%<br>Max Fil Eff: 99.50% | **1** | 85 | 8.3 | 0.158 | 0.500 | 99.50 |
|  | **2** | 85 | 8.0 | 0.477 | 1.19 | 98.81 |
|  | **3** | 85 | 7.7 | 0.404 | 0.910 | 99.09 |
|  | **4** | 85 | 7.8 | 0.464 | 0.865 | 99.14 |
|  | **5** | 85 | 8.1 | 0.256 | 0.736 | 99.26 |
|  | **Control 1** | 85 | 8.0 | 0.294 | 0.788 | 99.21 |
|  | **Control 2** | 85 | 8.1 | 0.294 | 0.803 | 99.20 |

Notes:
- The test method utilized in this assessment is not the NIOSH standard test procedure that is used for certification of respirators. Respirators assessed to this modified test plan do not necessarily meet the requirements of STP- 0059, and therefore cannot be considered equivalent to N95 respirators that were tested to STP-0059.

**Table 3.** Manikin Fit Evaluation.

| Manikin Fit Factor of Decontaminated N95s |  |  |  |  |  |
| --- | --- | --- | --- | --- | --- |
| Respirator Model, Decon Method, # of cycles | Treated Sample # | mFF Normal Breathing 1 | mFF Deep Breathing | mFF Normal Breathing 2 | Overall Manikin Fit Factor |
| 3M 1860, microwave plasma, 3 cycles<br>Static Advanced Medium Headform (Hanson Robotics) | 6 | 200+ | 200+ | 200+ | 200+ |
|  | 7 | 200+ | 200+ | 200+ | 200+ |
|  | Control 3 | 200+ | 200+ | 200+ | 200+ |
|  | Control 4 | 200+ | 200+ | 200+ | 200+ |

Notes:
- Per OSHA 1910.134\(f\)\(7\), if the fit factor as determined through an OSHA-accepted quantitative fit testing protocol is equal to or greater than 100 for tight-fitting half facepieces, then the fit test has been passed for that respirator. - This assessment does not include fit testing of people and only uses two exercises (normal and deep breathing) on a manikin headform. - This assessment is a laboratory evaluation using a manikin headform and varies greatly from the OSHA individual fit test. This headform testing only includes normal breathing and deep breathing on a stationary (non-moving) headform; therefore, fit results from this assessment cannot be directly translated to using the standard OSHA- accepted test. Instead, this testing provides an indication of the change in fit performance (if any) associated with the decontamination of respirators.

**Table 4.** Strap Integrity Evaluation.

| **Tensile Force in Respirator Straps of Decontaminated N95s**<br>**(recorded force values are at 150% strain)** |  |  |  |
| --- | --- | --- | --- |
| **Respirator Model, Decon**<br>**Method, # of cycles** | **Straps from Treated Sample #** | **Force in Top**<br>**Strap (N)** | **Force in Bottom**<br>**Strap (N)** |
| **3M 1860, microwave plasma, 3**<br>**cycles** | **1** | 2.925 | 2.814 |
|  | **2** | 2.894 | 2.846 |
|  | **3** | 2.974 | 2.837 |
|  | **Decontaminated Strap**<br>**Average** | 2.931 | 2.832 |
|  | **Control 1** | 2.839 | 2.921 |
|  | **Control 2** | 3.034 | 3.084 |
|  | **Control Strap Average** | 2.937 | 3.003 |
|  | **% Change**<br>**((Deconned - Controls) /**<br>**Controls)** | -0.2% | -5.69% |

Figure 3 shows an emission spectra taken during the bright glowing plasma using three different materials to make the antenna. The brightest plasmas were made using a copper antenna, though the coat-hanger (steel) antenna which was used for all of the decontamination experiments had all the same spectral features. The aluminum antenna produced the least bright plasmas and quickly melted after one or two uses.

**Figure 3:**
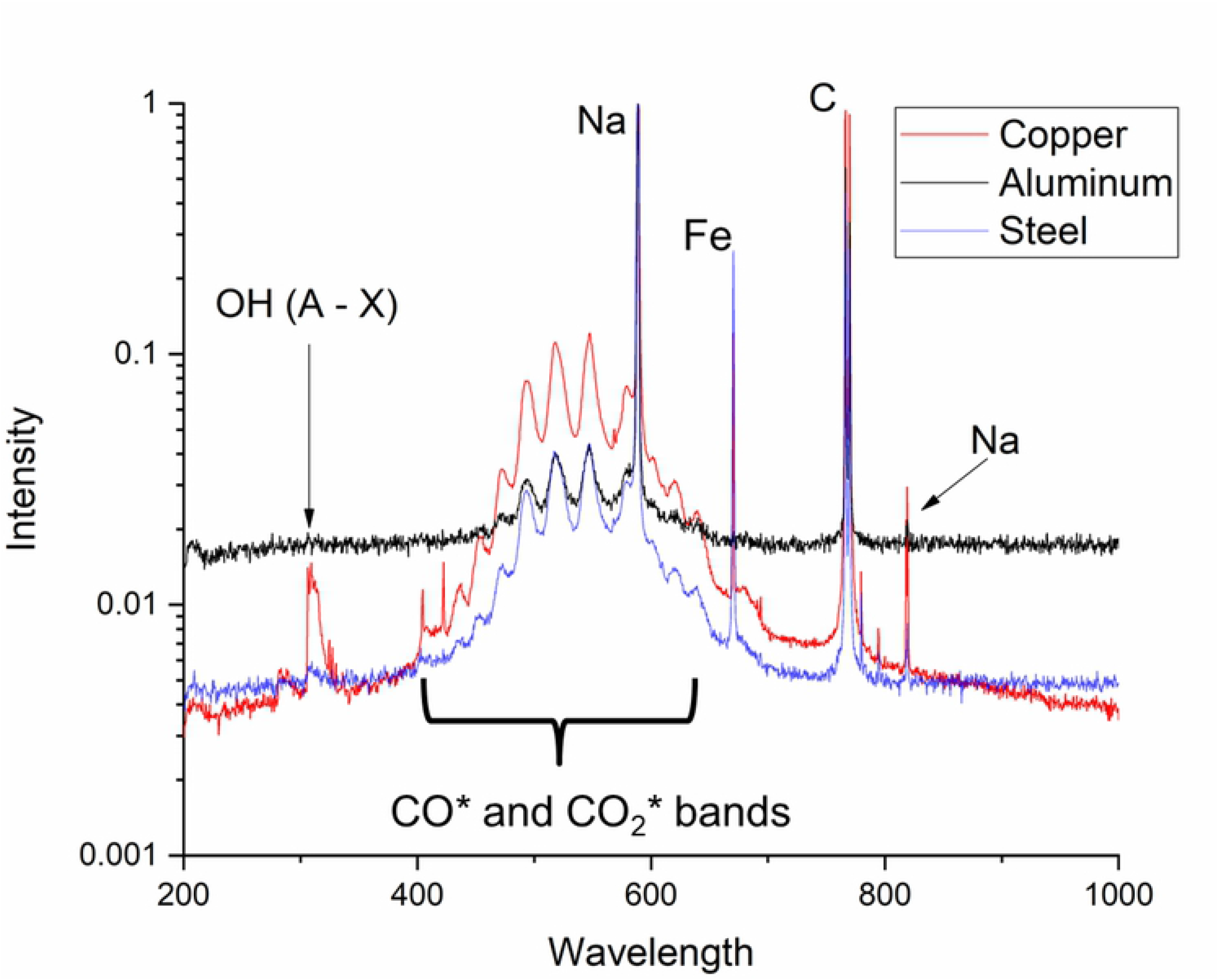
Spectra ofactive plasma region using three different antenna materials.

All of the spectra contained the Na line doublet at 589.0 and 589.6 nm and the C lines at 766.2 and 768.5 nm. The Cu and steel antenna showed an Fe line at 670.4 nm, but at a lower intensity. Of most interest is the molecular vibrational and rotational bands showing the presence of excited state radicals. The OH complex around 300nm is clearly seen in the brightest of plasmas. The continuum emission with a superimposed band structure between 400 and 650 nm is likely due to CO and CO_2_ excited states[9]. Every plasma heats the ceramic material under the gap to the point of becoming red hot. Since this is done in air, some combustion of the carbon-containing ceramic material is expected, and that may be the source for the carbon oxide compounds.

## IV. Discussion

A few notes of caution need to accompany the description given here. First, if possible, one should dispose of contaminated masks and use new ones to assure optimal safety. If this procedure is used to decontaminate N-95 respirators, it is important that an actual plasma is made. It is quite obvious when the system goes into the continuous mode: A bright light is emitted, and a loud buzzing sound is made by the plasma. The antenna gets very hot – so hot that the ends glow. The material beneath the antenna gap also gets very hot and may glow red. These surfaces should not be touched until they cool down. If plastic or other flammable material touches the red-hot glowing antenna or surface, it may catch on fire. This is likely why microwave manufacturers do not recommend that people put metal into microwave ovens. The metal gets hot, and if it is bent into the correct shape it can serve as an antenna and generate an atmospheric-pressure plasma as shown in this work. As shown by our results however, the plasma species generated are capable of decontamination in 30 seconds and anyone with a microwave oven and a few simple household items can convert it into a N-95 respirator decontamination unit for emergency use.

## Acknowledgements

Funding for this work was provided by the JUMP-ARCHES program of OSF Healthcare in conjunction with the University of Illinois. We’d like to thank Jeremy Neighbors from the office of occupational safety and health for providing the fit tests at Illinois, and Michael Bergman at the CDC’s NPPTL for the fit and filtration tests performed there.

## Author Bio

David Ruzic is the Able Bliss Professor in Nuclear, Plasma and Radiological Engineering and holds a zero-time appointment in the Carle-Illinois College of Medicine. He creates and studies plasma sources for use in microelectronic processing and fusion energy.

